# Single-cell microencapsulation improves lung retention of endothelial colony forming cells after intravascular delivery and unmasks therapeutic benefit in severe pulmonary arterial hypertension

**DOI:** 10.1101/2022.11.05.514522

**Authors:** Nicholas D. Cober, Ketul R. Chaudhary, Yupu Deng, Chyan-Jang Lee, Katelynn Rowe, David W. Courtman, Duncan J. Stewart

**Author notes:** **Corresponding author**: Duncan J Stewart., Executive Vice-President Research, The Ottawa Hospital; CEO & Scientific Director, Ottawa Hospital Research Institute; The Evelyne & Rowell Laishley Chair and Professor, Department of Medicine, University of Ottawa, 501 Smyth Road, Box/C.P. 511, Ottawa, ON, K1H 8L6. Both authors contributed equally to the work (Nicholas D. Cober, Ketul R. Chaudhary).

## Abstract

**Background:** Pulmonary arterial hypertension (PAH) is triggered by pulmonary vascular endothelial cell apoptosis and microvascular loss; therefore, therapies that can regenerate lost vasculature may offer therapeutic benefit. Endothelial colony forming cells (ECFCs) can directly repair damaged blood vessels and may have therapeutic potential for the treatment of PAH. However, poor retention of ECFCs in the lungs following intravenous delivery greatly limits their therapeutic application. Therefore, we studied whether cellular microencapsulation could enhance ECFCs viability and retention in the lung after systemic delivery and improve therapeutic efficacy of ECFCs in a rat monocrotaline (MCT) PAH model.

**Methods:** ECFCs were encapsulated by vortex-emulsion using various concentrations of agarose, and capsule size and initial cell viability were assessed. Encapsulated and free ECFCs were transduced with luciferase and administered to Sprague-Dawley rats three days after injection of MCT. ECFCs were tracked *in vivo* by bioluminescence imaging (BLI) to assess cell persistence and bio-distribution. At end-study, right ventricular systolic pressure (RVSP) and right ventricular hypertrophy were assessed for therapeutic efficacy.

**Results:** Microgel encapsulation using 3.5% agarose improved cells survival and supported cell migration from capsules. At 15 minutes after delivery, BLI radiance were similar for free and microencapsulated ECFCs; however, only encapsulated cells could be detected by BLI at 4 and 24 hours. Transplantation of microencapsulated ECFCs led to significant improvement in RVSP three weeks after delivery compared to non-encapsulated ECFCs.

**Conclusion:** Together, microencapsulation increased retention of ECFCs within the lungs. Furthermore, even a modest increase in ECFCs persistence over 24 hours can provide an important therapeutic benefit in the rat MCT model of PAH.

## Introduction

Cell-therapy has emerged as a promising option for treatment of various diseases and its potential is clearly evident by the ever-incising number of clinical trials using a wide variety cell types. Despite the tremendous promise that cell therapies offer for repairing and regenerating damaged tissues or organs, there has been only limited success in their translation into clinical therapies (1). A major factor in these mixed results is poor cell retention and survival of the transplanted cells. (2–5). Endothelial progenitor cells (EPCs) have been used to promote the regrowth and repair of damaged vasculature in many pre-clinical models of vascular diseases (6–8). These cells are classified as early-EPCs (E-EPCs or circulating angiogenic cells) or late-outgrowth EPCs (or endothelial colony forming cells; ECFCs) based on the time of appearance in culture (9). ECFCs are highly proliferative cells with an endothelial cell-like cobblestone morphology, and are capable of forming functional blood vessels (10). While E-EPCs act mainly by paracrine mechanisms, ECFCs are thought to participate directly in the repair and regeneration of blood vessels (11).

Pulmonary arterial hypertension (PAH) is a devastating lung vascular disease with 79% 3-year mortality (12). PAH is caused by damage and loss of the effective lung microvasculature, either by a degenerative mechanism or by occlusive arterial remodeling, leading to increased vascular resistance, as evidenced by increased pulmonary vascular resistance and pulmonary arterial pressures with eventual right ventricular failure (13,14). Current treatment options are limited, and available pharmacotherapies focus on targeting an imbalance between vaso-constrictor and -dilator factors within the pulmonary microcirculation (15,16). While current vasodilator therapies can improve symptoms and functional status, with the exception of parenteral prostacyclin their effects on survival are less certain and they are not curative. Since progressive lung arterial pruning is the fundamental pathological feature in PAH, a curative therapy would necessarily regenerate lost lung vasculature. In preclinical models of PAH, E-EPCs have been used to repair the lung microcirculation, with some improvements in pulmonary hemodynamics (4,6,17). While E-EPCs were effective in preventing PAH in a monocrotaline (MCT) induced rat model (6), ECFCs failed to improve pulmonary hemodynamics (4,5). We hypothesize that since ECFCs act directly by engraftment and incorporation into newly formed blood vessels, this lack of efficacy is most likely due to poor survival and retention after intravenous delivery.

Encapsulation of cells creates a protective microenvironment, which can improve transplanted cell survival and increase retention at the site of injury. Single cell microencapsulation provides a temporary micro-niche for the cells during transplantation and allows the delivery of cells into the circulation as a suspension. Microencapsulation of mesenchymal stromal cells within an agarose hydrogel increased survival of suspended cells, and improved cellular retention within a rat hind limb model (18). Microencapsulation of explant derived cardiac cells (EDCs) within nanoporous hydrogels promoted cell retention within the heart, reducing scar size and improving left ventricular ejection fraction post myocardial infarction (19,20).

Therefore, we studied whether microencapsulation of ECFCs would result in enhanced retention of ECFCs within the lungs, leading to increased therapeutic efficacy in an experimental model of PAH. We now demonstrate increased lung retention of microencapsulated ECFCs after intravascular delivery leading to the significant reduction on pulmonary arterial pressures and remodeling in the rat MCT model, which has not been previously demonstrated with ECFCs.

## Methods

### Endothelial cell isolation and culture

Rat bone marrow ECFCs and human L-EPCs were cultured and characterized (4). In brief, the mononuclear cell fraction of rat bone marrow or human peripheral blood was obtained by layering the cell suspension over Histopaque^®^-1083 (density: 1.083 g/mL, Sigma, ON, Canada) or Ficoll (density: 1.2 g/mL, Fisher Scientific, ON, Canada), respectively, followed by centrifugation at 400 g for 40 min. Mononuclear cells were collected from white layer at the interface of media (top) and Histopaque/Ficoll (bottom). Bone marrow mononuclear cells were plated in endothelial growth medium 2 + 10% fetal bovine serum (FBS) (EGM-2MV, Lonza, Switzerland) on fibronectin coated plates and cultured at 37°C in 5% CO_2_ incubators. Colonies with cells showing cobble stone-like endothelial cell morphology appeared in culture between 9 and 14 days after plating mononuclear cells. Endothelial-like colonies were isolated and further sub-cultured at 37°C in 5% CO_2_ incubators, passaging as necessary. In a separate procedure, all the cells cultured were passaged further without colony selection, and these cells were termed: culture modified bone-marrow cells (CM-BMCs). Otherwise, both ECFCs and CM-BMCs were processed similarly. Human umbilical vein endothelial cells (HUVEC, Lonza, Switzerland) were used in preliminary encapsulation experiments. HUVEC were cultured in endothelial growth medium 2 (EGM-2, Lonza, Switzerland) at 37°C in 5% CO_2_ incubator, passaging as necessary. Cells were used in experiments between P6 and P9.

### Flow cytometry

Rat ECFC or human L-EPCs were harvested and passed through 40 μm cell strainer to obtain single cell suspension (500,000/100 μL). Cells were incubated with primary antibodies (Table-1) for 30 min at 4 °C in dark. Following primary antibody, cells were washed 3 times in flow buffer followed by secondary antibody incubation for 30 min at 4 °C in dark. Cells were washed again for 3 times with flow buffer and then analyzed using Attune® Acoustic Focusing Flow Cytometer (ThermoFisher Scientific, ON, Canada), with analysis performed using FlowJo v 8.3 (FlowJo LLC, Ashland, OR, USA).

**Table-1.** Antibody source and concentration used for flow cytometry.

| **Target** | **Species** | **Source** | **Cat#** | **Dilution** |
| --- | --- | --- | --- | --- |
| vWF | Rat, human | Dako | A008202-5 | 1:400 |
| CD31 | Human | BD Pharmingen | 555446 | 1:100 |
|  | Rat | Novus Biologics | NB100-2284 | 1:50 |
| VEGFR2 | Human, Rat | Cell Signaling<br>Technology | 2479S | 1:100 |
| CD45 | Human | BD bioscience | 555482 | 1:5 |
|  | Rat | Biolegend | 202207 | 1:50 |
| CD34 | Human | BD-Pharmingen | 555822 | 1:5 |
|  | Rat | Santa Cruz | sc-7324 | 1:20 |
| VE-Cadherin | Rat | Santa Cruz | sc-6458 | 1:20 |
| CD146 | Rat | R & D | MAB3250 | 1:50 |
| CD14 | Human | BD-Pharmingen | 555398 | 1:5 |
| CD73 | Rat | BD-Pharmingen | 551123 | 1:50 |

### Acetylated low density lipoprotein (Ac-LDL) uptake and lectin binding

Rat ECFC or human L-EPCs (500,000/well) were plated on a 6 well plate and 24 hr later media was removed, cells were washed with PBS and incubated with DiI-Ac-LDL (10 µg/mL) for 3 hr at 37°C in 5% CO_2_ incubator. Following Ac-LDL incubation, cells were washed with PBS, harvested, and incubated with Fluorescein labeled Griffonia (Bandeiraea) Simplicifolia Lectin 1 (1:100) for 1 hr at room temperature. Following lectin binding cells were washed with PBS and fixed in 4% paraformaldehyde and analyzed by flow cytometry.

### Matrigel network formation assay

To confirm network forming ability of rat ECFC, cells were harvested and plated on growth factor-reduced Matrigel (Corning™ Matrigel™, Fisher Scientific, ON, Canada) in 96-well flat-bottom plate in EGM-2MV media and incubated for 16 hr at 37°C in 5% CO_2_ incubator. Images were captured using TS100 microscope (Nikon, ON, Canada).

### Lentiviral transduction

Rat ECFCs were transfected with a lentivirus construct under the murine stem cell virus (MSCV) promoter to overexpress red fluorescent protein (RFP) and luciferase. Fluorescently activated cell sorting was performed to isolate the RFP+ stably transfected cells, at the flow cytometry facility at the Ottawa Hospital Research Institute.

### Cellular encapsulation

ECFCs or HUVEC were re-suspended in PBS (Gibco, ON, Canada) with fibrinogen (0.05 mg/ml, Sigma Aldrich, ON, Canada), fibronectin (0.005 mg/ml, Roche, Switzerland) and Pluronic F-68 (0.33%, Sigma Aldrich, ON, Canada), prior to mixing with heated ultra-low gelling agarose type XI (Sigma Aldrich, ON, Canada). The well mixed cell-hydrogel suspension was added dropwise to warm dimethylpolysiloxane (PDMS, Sigma Aldrich, ON, Canada), vortexed for 1 minute, and placed in an ice bath for 10 minutes. As previously described (18), encapsulated cells were washed with Hank’s balanced salt solution (HBSS, Gibco, ON, Canada) to remove PDMS phase, and filtered to establish the encapsulated cell population.

### Assessment of cell viability of encapsulated cells

Encapsulated cells were assessed *in vitro* for viability by WST-1 assay and Calcein AM staining. Encapsulated or adherent cells were plated in a 96 well plate for 24 or 48h when WST-1 reagent (Roche, Switzerland) was added, or Calcein AM (Invitrogen, ON, Canada) and Nuclear Blue live ready reagent (Invitrogen, ON, Canada) were added. WST-1 assay was assessed by absorbance after 2 hours with an Omega PolarStar (BMG Labtech, Germany). Live imaging was performed using a Zeiss Observer Z1 inverted microscope (Zeiss, Germany) and images were quantified using FIJI for ratio of calcein positive cells to total counted nuclei.

### Assessment of cell migration from agarose microcapsules

Migration of encapsulated cells was performed with a Boyden chamber assay in a 24 well plate as per manufacturer’s recommendations. Briefly, encapsulated cells were plated in basal media on the top and complete media containing 20% FBS was added to the bottom well. Cells were incubated for 24h. Cells were stained with Nuclear Blue live ready reagent (Invitrogen, ON, Canada) and images were acquired with the Zeiss Observer Z1 inverted microscope (Zeiss, Germany). Images were quantified as before using FIJI.

### Monocrotaline model of PAH and cell injections and hemodynamic measurements

All animal experiments were approved by the University of Ottawa Animal Care Committee and conducted according to the guidelines from the Canadian Council for Animal Care. Female Sprague Dawley (SD) rats (150 – 175g; Charles River) were injected with monocrotaline (MCT) (50mg/kg, Sigma Aldrich, ON, Canada) by intra peritoneal (i.p.) injection three days before cell injections. For cell injections, rats were anaesthetized using isoflurane inhalation (induction: 5% isoflurane 1 L/min; maintenance: 2% isoflurane 1 L/min), right jugular vein was isolated, and the angiocath was inserted into the jugular vein. Each animal received 1,000,000 cells or vehicle (0.5 mL PBS). Following injection, topical bupivacaine was applied immediately after wound closure and twice daily for one day post-surgery. Buprenorphine (s.c., 0.03mg/kg) was administered 1 hour prior to surgery and once daily for two days post surgery for pain management. At 24 days post MCT animals were anaesthetized by an i.p. injection of ketamine (35 mg/kg) and xyalzine (7 mg/kg) and the right ventricular systolic pressure (RVSP) was measured, as previously described (6,21) The lungs and heart were harvested for histology and protein, and the ratio of right ventricle (RV) to left ventricle plus septum (LV+S) was measured, as previously described (6,21)

### Cell retention analysis by qPCR

To study cell retention, we injected male cells into female rats and assessed cell retention using qPCR for sex-determining region of Y chromosome (Sry) gene. Briefly, female rats were subjected to MCT model and on day-3 male ECFCs were injected. Samples were collected at 15 min and 21 days after ECFCs injection. DNA was extracted from lung samples and 5,000,000 male ECFCs using DNeasy Blood & Tissue Kit (Qiagen, ON, Canada) and quantified using Quant-iT™ PicoGreen™ dsDNA Assay Kits (ThermoFisher Scientific, ON, Canada). Standard curve of different ratios of male ECFC DNA to female rat lung DNA was prepared (1, 0.1, 0.001, 0.0001 ng/1000 ng total DNA) and final sample DNA concentration was adjusted to 125ng DNA/μL. qPCR analysis of standards and samples was performed using QuantiTect SYBR Green PCR Kit (Qiagen, ON, Canada) as per manufacturer’s recommendation using primers for amplifying Sry gene (Sry Forward-AAGTCAAGCGCCCCATGA, Sry reverse-TGAGCCAACTTGTGCCTCTCT). Amount of male DNA in total DNA of lung samples was calculated from standard curve and used to quantify cell retention.

### Assessment of the effect of encapsulated ECFCs in MCT model

SD rats were treated with MCT and, ECFCs (300,000 cells in 0.5 mL PBS), encapsulated-ECFCs (300,000 cells in 0.5 mL PBS) or empty capsules (equivalent to 300,000 encapsulated cells in 0.5 mL PBS) were injected through jugular vein as described above. The dose of capsulated cells was lowered as higher dose of capsules were not tolerated by MCT treated rats.

### *In vivo* bioluminescent imaging

At 0, 4 and 24 hr post cell injection (**Supplemental Figure 1**), animals were injected with D-luciferin (150 mg/kg, ip; Biovision, CA, USA). These animals were subsequently anaesthetized by isoflurane and imaged using the IVIS spectrum (PerkinElmer, MA, USA) at the University of Ottawa Preclinical Imaging Core (RRID:SCR_021832). Animals were euthanized before subsequent open chest and *ex vivo* lung images were taken to account for the thickness of the rat’s tissue. Bioluminescent imaging (BLI) was quantified using Living Image 2.0 software (PerkinElmer, MA, USA).

### Immunohistochemistry

As previously described (21), lungs were inflated with 1:1 ratio of OCT:Saline and fixed in 4% paraformaldehyde solution (Electron Microscopy Science, PA, USA) before dehydration and paraffin embedding. Sections of fixed lung tissue (5µm thickness) were cut and stained with hematoxylin (Vector Laboratories, CA, USA) and eosin (ThermoFisher Scientific, ON, Canada) using standard protocol. Alternatively, sections were blocked with 2% goat serum (Rockland Immunochemicals, PA, USA) and stained with primary rabbit anti-firefly luciferase antibody (Abcam, Cambridge, UK) 1:100 overnight at 4°C followed by Alexa Fluor 594 goat-anti rabbit secondary antibodies (Invitrogen, ON, Canada) at 1:400 for 1 hr at room temperature. Immunofluorescent images were analyzed with a Zeiss M2 (Zeiss, Germany) and prepared with FIJI. Analysis of vascularization was done by grading blood vessels that were non-muscularized (0), partially muscularized (1), or fully muscularized (2), representative images shown in **Supplemental Figure 2**, as previously performed (4). All vessels were counted and assessed in one lung section per animal by a blinded reviewer.

### Statistical Analysis

All data are presented as means ± SEM. Differences between groups were analyzed by one-way analysis of variance (ANOVA), with appropriate post-hoc comparisons. A p-value of p<0.05 was considered significant. All statistical analysis was performed with Graph Pad Prism 7.0 (Graph Pad, CA, USA).

## Results

### Microgel encapsulation supports endothelial cell survival

The vortex emulsion method was optimized using HUVECs and we compared cells encapsulated in 2 to 5% agarose capsules supplemented with or without fibronectin and fibrinogen (**Figure 1A**). The resulting microcapsules were heterogeneous in size, ranging from 10 – 70 µm in diameter. Higher concentrations of agarose (5%) produced larger capsules compared to lower concentrations of agarose (2%) (**Figure 1B**). Next, we evaluated the effect of agarose concentration on cell survival and egress from the capsule as these are important characteristics for the effectiveness of cell therapy. We observed decreased cell egress over 24 hours from capsules with increasing agarose concentration (**Figure 1C**). These data suggest that the stiffer hydrogels delayed the egress of cells. The concentration of agarose had no significant impact on cell survival under normal condition, as assessed by WST-1 assay and CalceinAM staining (**Figure 1D and E**, respectively). However, in response to serum starvation, conditions that better reflect those that cells are exposed to during cell transport and delivery, cells encapsulated in 3.5 and 5% agarose exhibited superior survival compared to adherent cells (28% vs 71% reduction, respectively) (**Figure 1F**). Based on these results and previous work (19), 3.5% agarose capsules were used for the *in vivo* studies.

**Figure 1:**
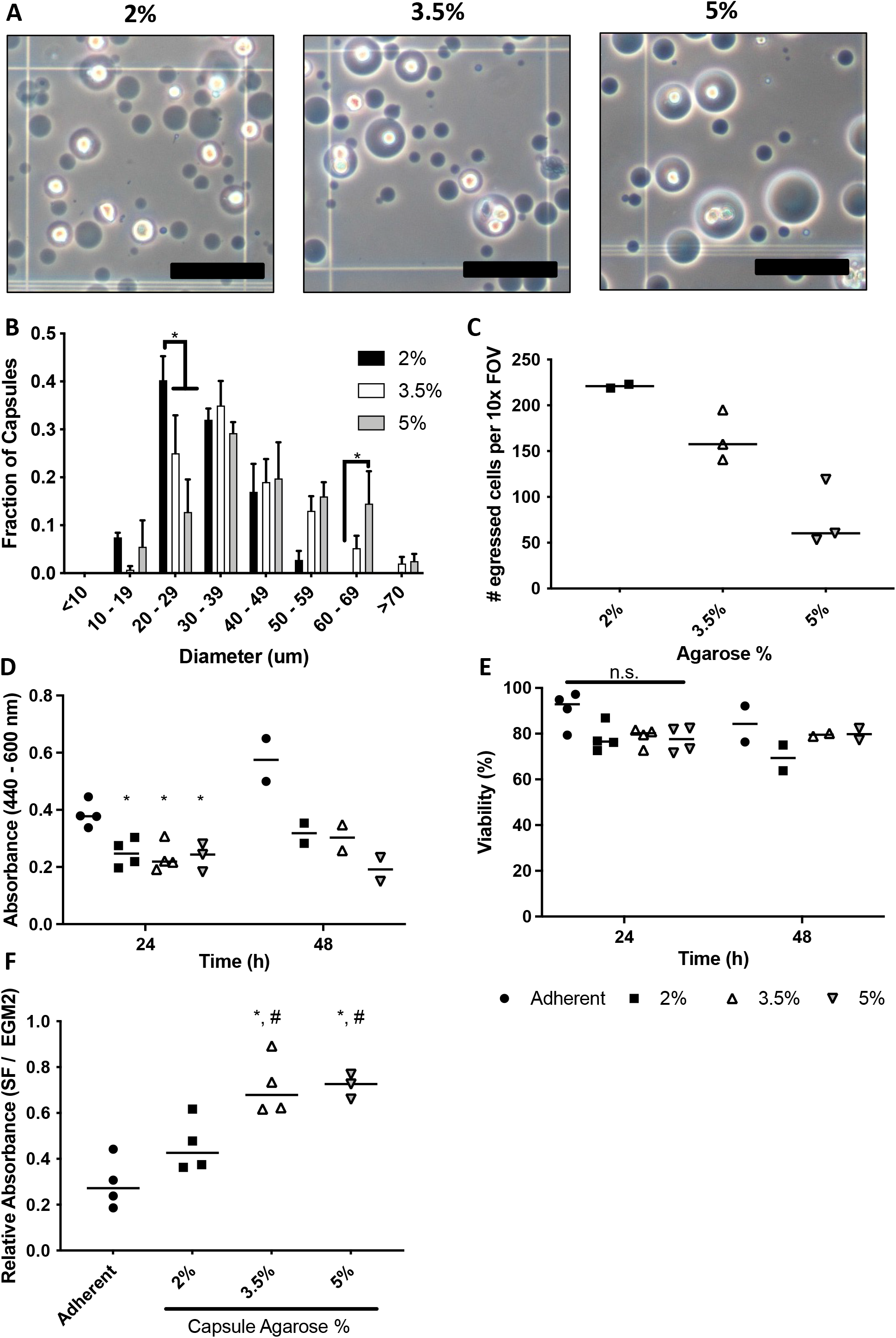
Varied agarose concentration has minimal effect on cell viability. (A) Representative phase contrast images of encapsulated cells in 2-5% agarose microgels, scale bars represent 100 µm. (B) Capsules containing cells appear to increase in diameter with increasing concentration of agarose in the hydrogel mixture. (C) Higher concentrations of agarose capsules appears to slow migration of cells from the capsules. Viability was unaffected by agarose capsule concentration as assessed by (D) WST-1 assay and (E) CalceinAM staining. Capsules provided a protective niche during 24h culture in serum free condition, shown by (F) raw WST-1 absorbance and (G) reduction in absorbance. Scale bar represents 100µm, * represents p < 0.05 vs adherent, # represents p < 0.05 vs 2% agarose. Data represents means ± SEM, n = 2-4.

### ECFCs were ineffective in severe pulmonary arterial hypertension

Rat ECFCs were isolated from the SD rat bone marrow and displayed typical endothelial growth characteristics. Cell morphology and surface markers of ECFCs were compared to human L-EPCs and CM-BMCs (**Figure 2)**. ECFCs displayed a cobblestone morphology when cultured on plastic similar to that of L-EPCs and HUVECs (**Figure 2A**) whereas CM-BMC exhibited more mesenchymal cell morphology. ECFCs demonstrated significant proliferative potential *in vitro* (**Supplemental Figure 3**). Furthermore, human L-EPCs and rat ECFCs shared common endothelial characteristics including uptake of ac-LDL, lectin binding, and expression of vWF and CD31. Rat ECFCs additionally expressed vascular endothelial cadherin, and low levels of vascular endothelial growth receptor 2 (VEGFR2), CD146, and CD34 (**Figure 2B**). Moreover, human L-EPCs and rat ECFCs completely lacked markers of leukocyte or monocyte cell lineage such as CD45 and CD14 (**Figure 2B**). Importantly, while CM-BMCs did exhibit some endothelial features (acLDL uptake, lectin binding and low vWF expression), they expressed the mesenchymal marker, CD73. These data confirm the endothelial identity of ECFCs and suggest that selection of cells is required to avoid contamination of mesenchymal-like cells from bone marrow.

**Figure 2:**
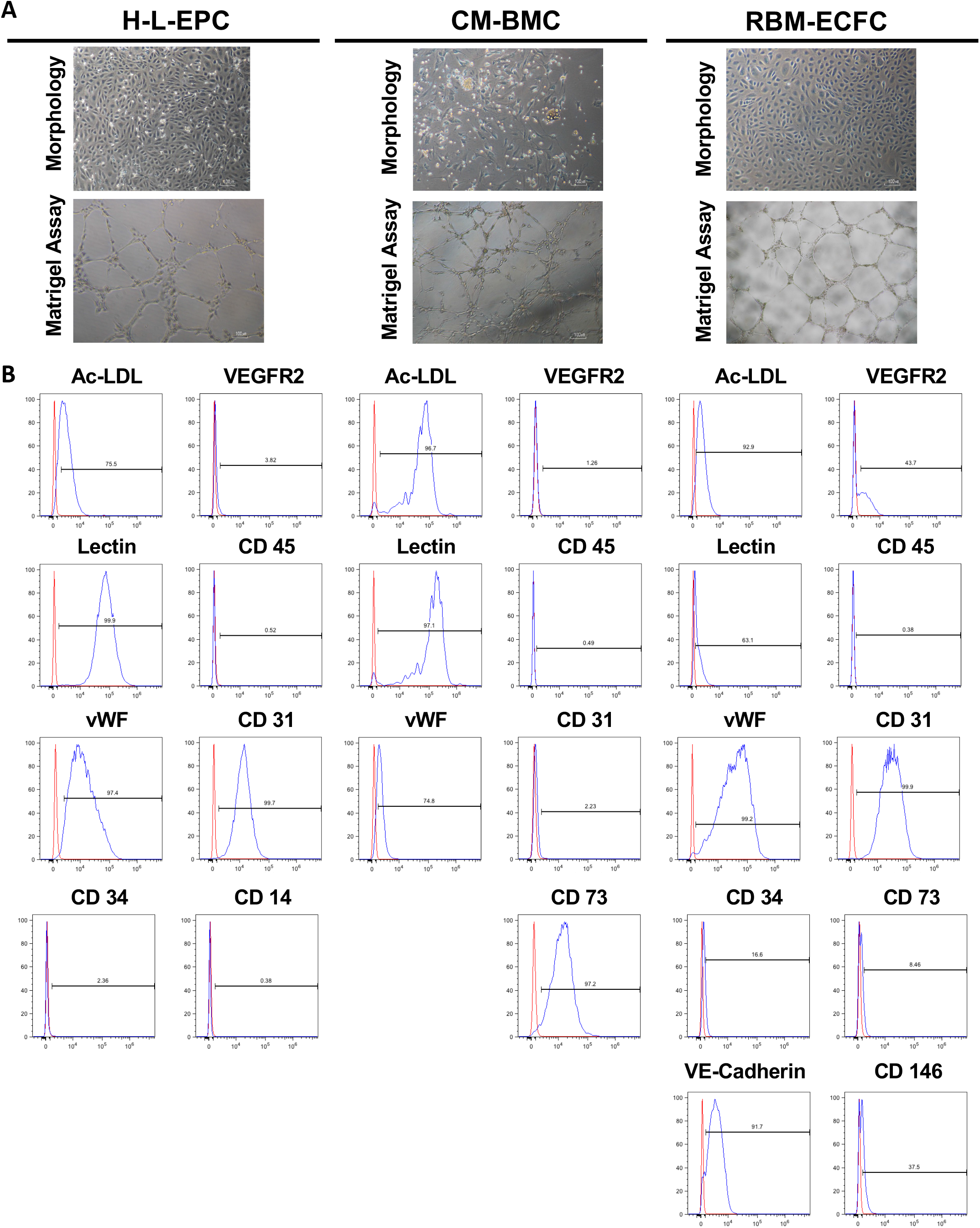
Rat ECFCs share morphology and surface markers with human L-EPCs. (A) Representative images of human L-EPC, rat CM-BMC, and rat ECFCs demonstrating the cobblestone morphology commonly observed in cultured endothelial cells. (B) Surface marker characterization by flow cytometry demonstrating the similarities between human and rat derived L-EPCs positive for ac-LDL, lectin, von Willebrand Factor, and CD31, while negative for CD45. The red line depicts isotype control and blue line represents staining with specific marker.

Next, we assessed the therapeutic efficacy of ECFCs in the MCT model of PAH. ECFC were delivered by intrajugular vein injection (1,000,000 cells) 3 days after MCT. Compared with vehicle control (PBS), treatment with ECFCs resulted in no detectable improvements in RVSP or RV remodeling at day 24, 3 weeks after cell delivery (**Supplemental Figure 4**). We assessed retention of the cells in the lungs by qPCR. At 15 minutes, we observed 80% retention of cells in the lungs; however, at the end study, cells were not detectable suggesting poor retention of syngeneic ECFCs following injection (**Supplemental Figure 4**).

### Encapsulation of ECFCs increased their retention within the lungs of MCT treated rats

To enhance the survival and retention of ECFCs, we performed microencapsulation in 3.5% agarose. A dose of 300,000 ECFC loaded microgels were injected by intrajugular vein injection. Bioluminescent imaging allowed tracking of live cells containing luciferase luminescent reporter.

The baseline signal within 30 minutes of cell injection was comparable between encapsulated and non-encapsulated ECFC treated animals; however, the signal for non-encapsulated cells was rapidly lost within 4 hours of cell injection and there was no detectable signal with either live or *ex vivo* imaging (**Figure 3A and B**). In contrast, encapsulated ECFCs could be detected up to 24 h post cell injection (**Figure 3A and B**), but not after 72 h even with *ex vivo* imaging. The retention of encapsulated cells was further verified by H&E and immunohistochemical staining with anti-luciferase antibody at 4h and up to 72h (**Figure 3C and D**). Additionally, empty capsules were found to persist *in vivo* for up to 21 days (**Supplemental Figure 5**). Therefore, encapsulation of ECFCs lead to a significant prolongation of lung retention in monocrotaline-treated rats.

**Figure 3:**
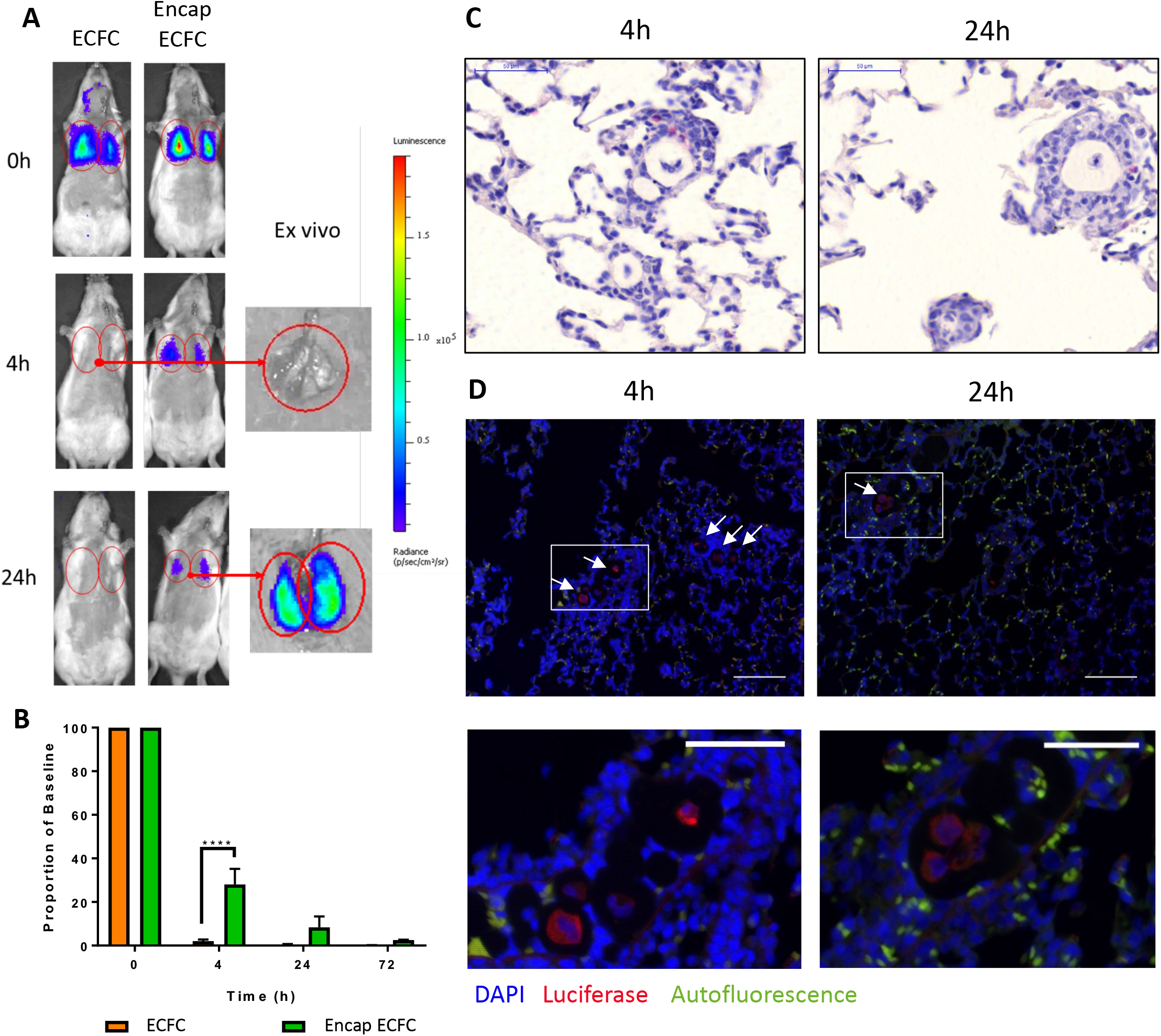
Encapsulated ECFCs are retained significantly longer than non-encapsulated cells. (A) Representative images of the bioluminescent signal emitted by luciferase transfected ECFCs 15 minutes after luciferin injection, including *ex vivo* images of lungs containing encapsulated cells. The encapsulated cells were detectable in the lungs up to 24h whereas non-encapsulated cells were cleared after 4h and not detectable even with *ex vivo* imaging. (B) The proportion of the baseline signal demonstrates at 24h ∼10% of the encapsulated cells were detectable by bioluminescent imaging. Representative images of encapsulated cells with H&E (C) (scale bar – 50um) or stained with anti-luciferase antibody (red), DAPI (blue), and auto fluorescence overlap (green) (D) at 4 and 24h post injection (scale bar – 100µm, or 50µm for zoomed image, demonstrating persistence of encapsulated cells *in vivo*. Data represented as mean ± SD, n =3-4.

### Encapsulated ECFCs prevented the onset of severe PAH and improved lung vascular morphology

Animals were followed for 24 days post MCT injection to assess the effect of cell delivery of pulmonary hemodynamics and vascular remodelling. No significant survival difference between ECFC treated rats and PBS treated rats was observed. Survival was also similar for rats treated with empty capsules or encapsulated ECFCs. Importantly, a trend towards lower survival in the groups treated with capsule was evident compared to the groups receiving capsules (empty capsules compared to PBS and encapsulated ECFCs compared to ECFCs). The only significant difference in survival was observed between ECFCs treated rats and empty capsules (**Figure 4A**). Again, animals treated with non-encapsulated ECFCs showed no improvement in RVSP or RVH compared to control-vehicle treated animals (**Figure 4B and C**). However, animals receiving encapsulated ECFCs demonstrated a significant improvement in RVSP (p < 0.05) compared to rats treated with vehicle control or empty capsules (**Figure 4B**). MCT-induced pulmonary hypertension was associated with arterial muscularization with an increase in the ratio of partially (33.8%) and fully (60.3%) muscularized vessels (**Figure 4D and E**). Non-encapsulated ECFCs or empty capsules had no effect on the severity of arterial muscularization. In contrast, treatment with encapsulated ECFCs significantly reduced muscularization (**Figure 4D**), yet no changes in overall lung vasculature were observed by MicroCT (**Supplemental Figure 6**). Taken together, these data suggest that the modest increase in retention of ECFCs resulting from microencapsulation was sufficient to uncover a therapeutic benefit in a model of PAH.

**Figure 4:**
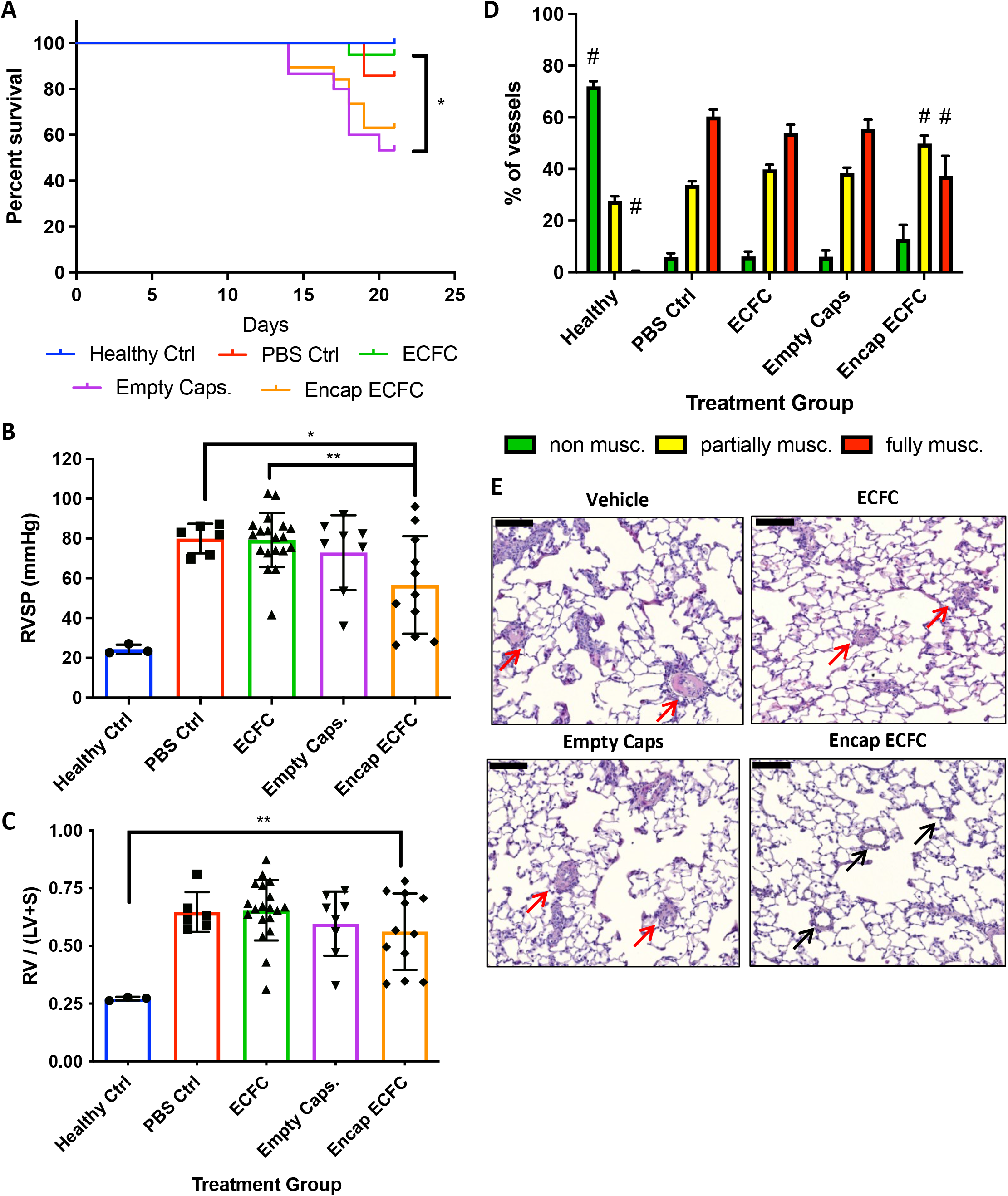
Capsules may be detrimental in animal survival, yet encapsulated ECFCs provided a therapeutic benefit to preventing development of severe PAH. (A) Survival curve demonstrating a slight detrimental effect of capsules on animal survival. (B) RVSP and (C) RVH for all surviving animals demonstrate the beneficial effect offered by encapsulated ECFCas in preventing the onset of severe PAH. (D) Severity of muscularization was graded and assessed to be (0) non-muscularized, (1) partially muscularized, or (2) fully muscularized, where animals treated with encapsulated ECFCs had fewer fully muscularized blood vessels, # represents p < 0.01 vs PBS ctrl. (E) Representative images of H&E stained histological sections demonstrating severe muscularization of blood vessels (red arrows) 24 days post MCT in animals treated with PBS vehicle, ECFCs, and empty capsules, whereas animals treated with encapsulated ECFCs had improved vascular remodelling (black arrows). Scale bar represents 100µm. Data represented as means ± SEM, n = 3 – 20.

## Discussion

ECFCs have tremendous promise for the treatment of vascular diseases due to their exceptional proliferative and angiogenic potential. ECFCs have been successfully used to repair and regenerate vasculature in many cardiovascular diseases (22) Therefore, it is surprising that ECFCs have failed to be effective in preclinical models of PAH (4,5), despite the critical role of arterial pruning in its pathogenesis (13). However, unlike E-EPCs which act nearly exclusively by paracrine mechanisms, ECFCs (or L-EPCs) are thought to act directly to regenerate blood vessels by engraftment and differentiation (8). Thus, the lack of cell persistence in the lung after intra venous delivery may be a major limitation to their potential repair of the lung arterial bed through direct integration. Therefore, we investigated whether a strategy to enhance the retention of ECFCs within the lungs would enhance their therapeutic potential using single cell microencapsulation.

Microencapsulation of cells within agarose hydrogels supplemented with fibrinogen and fibronectin offers a protective niche for the cells during transplantation. Similar strategies have been employed with MSCs (18), and EDCs for myocardial infarction (19,20). By incorporating integrin binding proteins it creates a protective niche within the hydrogel preventing anoikis and enhancing cell survival (18). In this report, we investigated the impact of hydrogel concentration for ECs encapsulation, to maintain cell viability under serum starved conditions, while allowing cell migration from capsules. Unlike macro-encapsulation that aims to permanently isolate cells from the host immune cells (23), single-cell microencapsulation provides a temporary shield to protect the cells during the traumatic effects of transplantation, yet allows cell egress within the first few days. The size of the cell-loaded microgels allows for ease of delivery by intravenous injection. There is also evidence that microencapsulation can promote increased paracrine secretions compared in 2D cell culture or bulk encapsulation systems (24) providing additional benefits to this delivery system.

Moreover, microencapsulation of ECFCs improved local retention of cells within the lungs, since even the smallest capsules were too large to traverse through the lung capillary bed and most would be retained within pre-capillary arterioles. In contrast, free ECFCs were rapidly cleared from the lungs within 4h of injection. We observed that the increase in retention of microencapsulated ECFCs within the damaged lung unmasked a therapeutic effect in the MCT model of PAH that was not evident with cells alone. Considering that the dose of microencapsulated ECFCs was only 300,000, compared to previous studies delivering up to 1.5 million free cells (4), this strongly supports the critical importance of ECFC retention for enhancing therapeutic effects. Although, the capsules provided short term improvement in ECFC retention, there was no evidence of cell persistence beyond 72 hours, whereas empty capsules could be found in the lung for at least three weeks post injection. This suggests that cells did not remain in capsules indefinitely, and either egressed from the capsules or underwent apoptosis after several days. However, even if a substantial proportion of cells exited the capsules, they did not persistent in the lung in sufficient numbers to be detectable by bioluminescent imaging, implying that the observed benefit may have involved paracrine mechanisms. Indeed, microencapsulation has previously been shown to enhance the paracrine effects of both MSCs and EDCs (19,24).

There are some limitations of our study. Despite the ease of encapsulation using the vortex-emulsion method, the capsules produced are heterogenous in size and many do not contain cells. Additionally, the current capsule formulation persists within the lungs for prolonged periods. The persistence of these capsules *in vivo*, and the variability in size, likely contributes to a weak trend toward worsening survival. It is possible that this could be due to a detrimental effect of the agarose hydrogel. For example, capsules with diameters above 50µm could block larger arterioles, thereby aggravating the underlying pruning of the arteriolar bed seen in this model, possibly explaining why there was no improvement in lung arterial architecture by micro-CT. This might be avoided by using a microfluidic technique to generate more homogeneous capsules and reduce the number of empty capsules (24,25). Reduction in capsule size heterogeneity, ratio of encapsulated cells to empty capsules, and degradability may allow larger doses of capsules to further enhance the therapeutic efficacy of this technology. Furthermore, optimization of the biomaterial composition could reduce the inflammatory response and allow enhanced degradability of the matrix for use in pulmonary vascular regeneration (26).

This study represents a proof-of-principle for the potential of microencapsulation technology to enhance the efficacy of ECFCs for treatment of PAH. Our microencapsulation strategy increased lung retention of ECFCs compared to the non-encapsulated controls. Furthermore, short term increases in retention resulted in hemodynamic improvement in the MCT model of PAH, although there is room to improve this approach by further optimizing the consistency and biocompatibility of the hydrogel capsules.

