## Supplemental Figures + Legends for "Single-cell microencapsulation improves lung retention of endothelial colony forming cells after intravascular delivery and unmasks therapeutic benefit in severe pulmonary arterial hypertension"

### Supplemental Figure Legends

**Supplemental Figure 1: Study design and treatment groups.** (A) Animals were i.p. injected with MCT, and provided preventive cell therapy 3 days post MCT. Bioluminescent imaging were done at baseline (0h), 4h, 1, 3 and 21 days post cell injection. End study at 24 days post MCT was performed to assess RVSP, RVH, and collect samples for histology. (B) Treatment groups given 3 days post MCT.

**Supplementary Figure 2. Examples of blood vessels used for grading vascular muscularization**. Grade 0 indicates no muscularization of blood vessel, Grade 1 indicates minor or partial muscularization of the blood vessel has occurred, grade 2 indicates a severely muscularized blood vessel where only a partial lumen is observed.

**Supplemental Figure 3: Characterization of L-EPCs proliferative potential.**

**Supplemental Figure 4: Lack of retention within MCT treated lungs after 21 days.** Green rats were sacrificed before 21 days post cell injection due to reaching humane end-point.

**Supplemental Figure 5: Capsule degradation.** (A) Representative images of capsules within the lungs after 4, 24, 72 hours or 21 days post cell injection, indicated with black arrows. (B) Average number of capsules counted per mm2 of lung indicating capsules are cleared from the lung over time. (C) Capsule diameter as assessed over time, despite fewer capsules present by 21 days the capsules remaining are not significantly changed in size. n = 3 per time point, data represented as mean ± SD.

**Supplemental Figure 6: Right ventricular systolic pressure strongly correlates with vascular volume within the lungs.** (A) Representative images of MicroCT analysis from the left lobes of SD rats given MCT and treated with: empty capsules, L-EPCs, or encapsulated L-EPCs. (B) Vascular volume between treatment groups compared by vessel diameter suggesting this experiment was underpowered to identify differences between these groups. (C) RVSP and total vascular volume had a strong inverse relationship suggesting that high pressures are more likely when total volume is decreased. Data represented as means ± SEM, n = 3 – 4.

Sup Fig 1

A

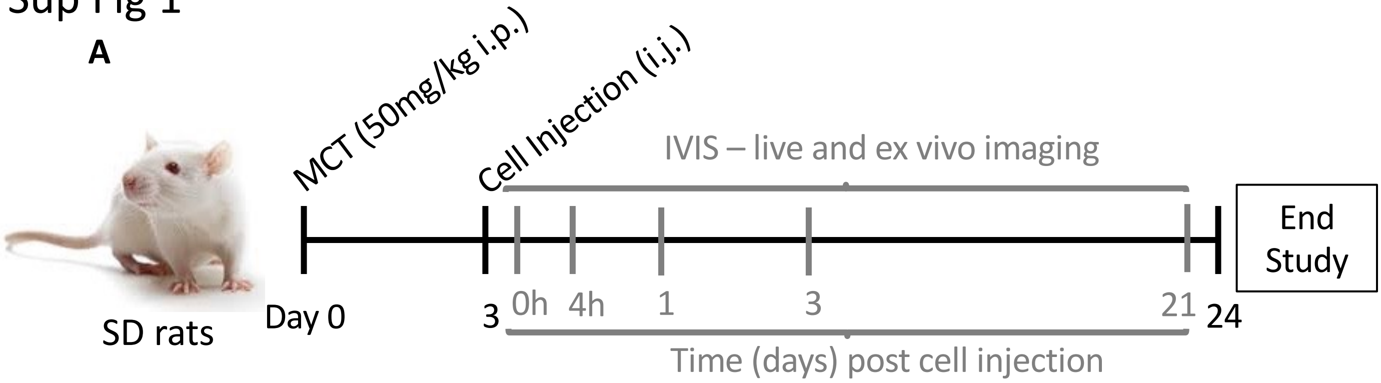


B

| Treatment Group | MCT | L-EPC | Capsules |
|---|---|---|---|
| Healthy Control | - | - | - |
| PBS Vehicle Control | + | - | - |
| Non-encapsulated ECFC | + | + | - |
| Encapsulated ECFC | + | + | + |
| Empty Capsules | + | - | + |

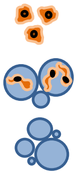

Sup Fig 2

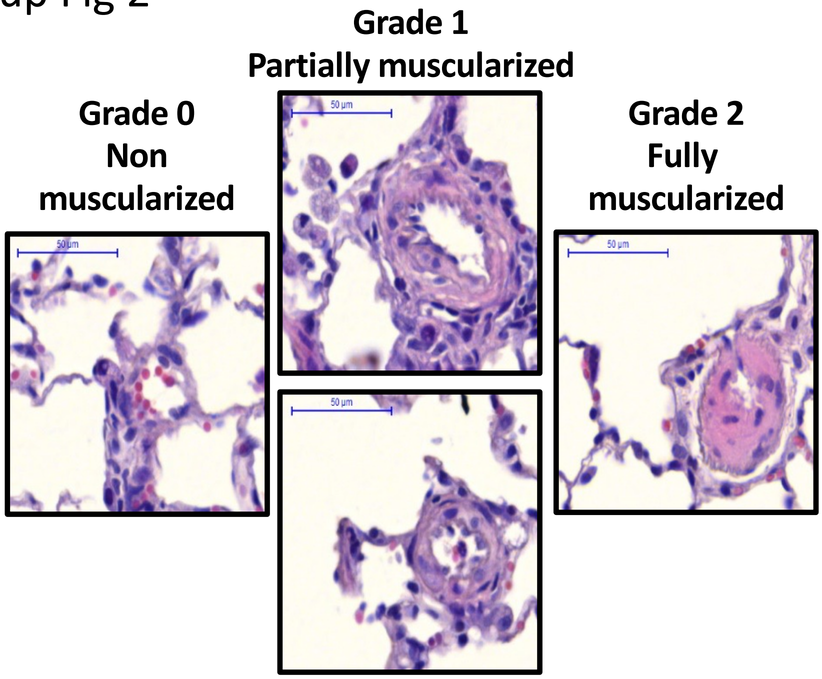
Grade 1
Partially muscularized
Grade 0
Non
muscularized
Grade 2
Fully
muscularized
50 µm
50 µm
50 µm
50 µm

Sup Fig 3

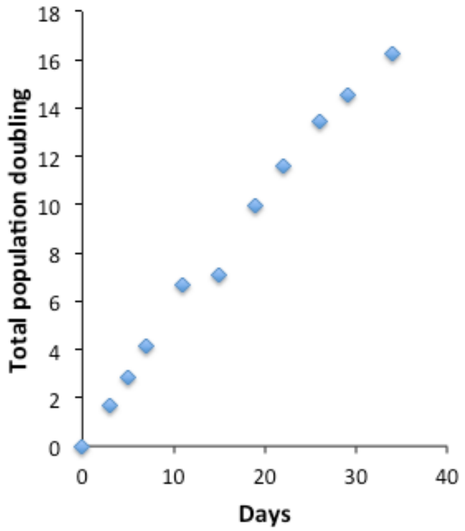

Total population doubling
18
16
14
12
10
8
6
4
2
0
0
10
20
30
40
Days

Sup Fig 4

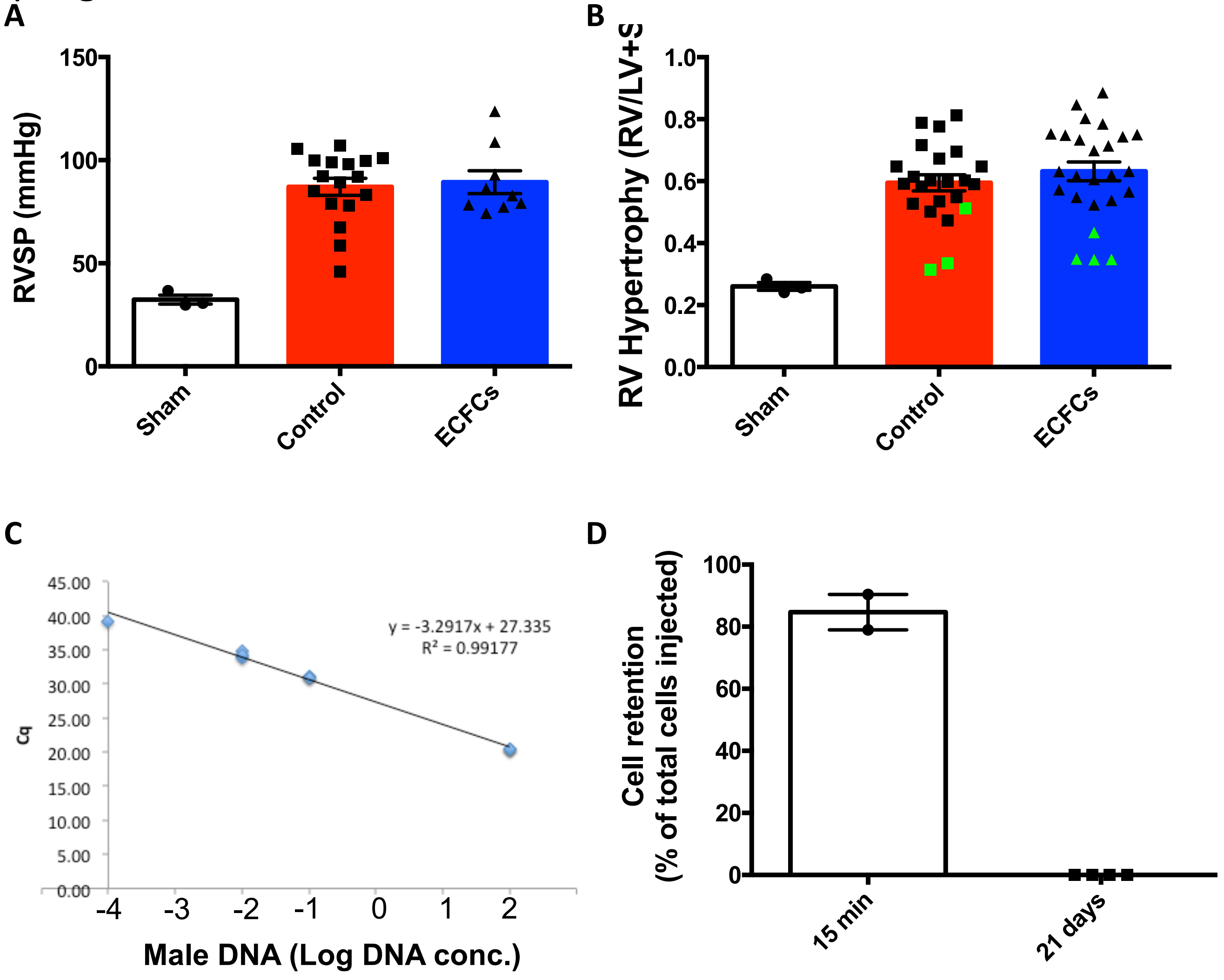
A
RVSP (mmHg)
150
100
50
0
Sham
Control
ECFCs
B
RV Hypertrophy (RV/LV+S
1.0
0.8
0.6
0.4
0.2
0.0
Sham
Control
ECFCs
C
45.00
40.00
35.00
30.00
25.00
20.00
15.00
10.00
5.00
0.00
Cq
y = -3.2917x + 27.335
R² = 0.99177
-4
-3
-2
-1
0
1
2
Male DNA (Log DNA conc.)
D
Cell retention
(% of total cells injected)
100
80
60
40
20
0
15 min
21 days

Sup Fig 5

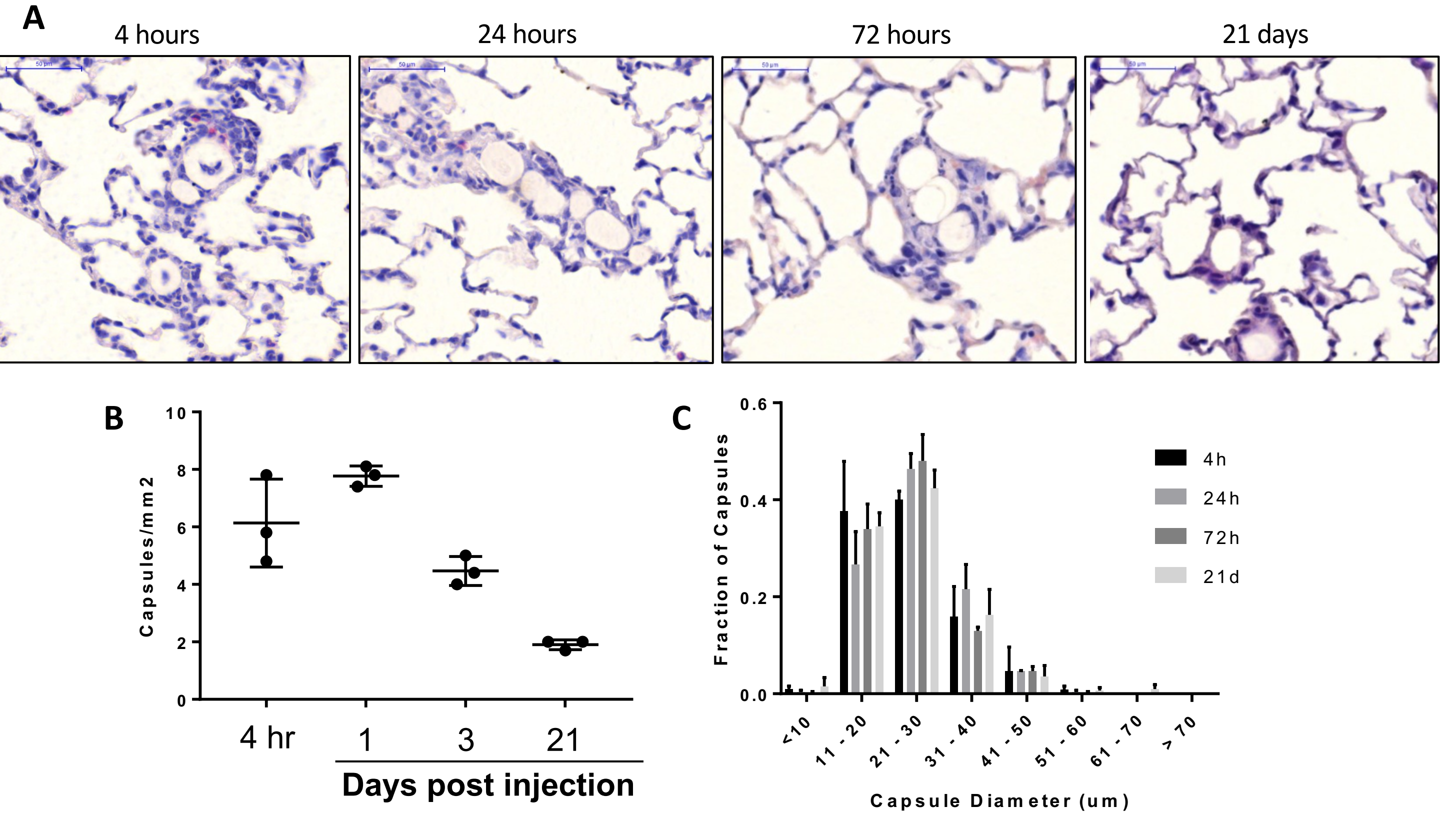
A
4 hours
24 hours
72 hours
21 days
B
Capsules/mm2
0
2
4
6
8
10
4 hr
1
3
21
Days post injection
C
Fraction of Capsules
0.0
0.2
0.4
0.6
4h
24h
72h
21d
<10
11 - 20
21 - 30
31 - 40
41 - 50
51 - 60
61 - 70
> 70
Capsule Diameter (um)

Sup Fig 6

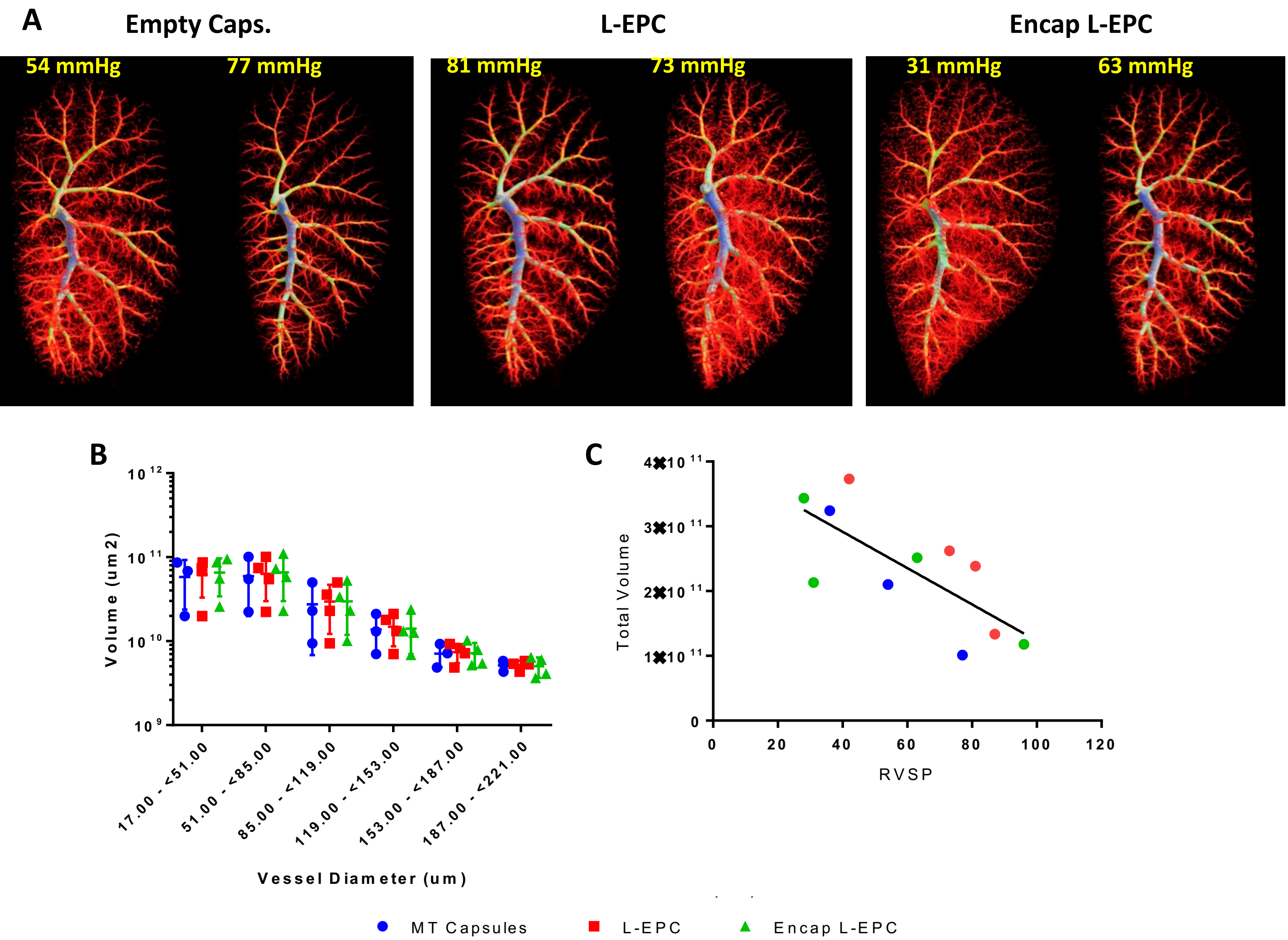
A
Empty Caps.
L-EPC
Encap L-EPC
54 mmHg
77 mmHg
81 mmHg
73 mmHg
31 mmHg
63 mmHg
B
Volume (um2)
10^12
10^11
10^10
10^9
17.00 - <51.00
51.00 - <85.00
85.00 - <119.00
119.00 - <153.00
153.00 - <187.00
187.00 - <221.00
Vessel Diameter (um)
C
Total Volume
4×10^11
3×10^11
2×10^11
1×10^11
0
0
20
40
60
80
100
120
RVSP
MT Capsules
L-EPC
Encap L-EPC
